# Widespread dynamic and pleiotropic expression of the melanocortin-1-receptor (MC1R) system is conserved across chick, mouse and human embryonic development

**DOI:** 10.1101/212712

**Authors:** Anna C. Thomas, Pauline Heux, Chloe Santos, Wisenave Arulvasan, Nita Solanky, Magalie E. Carey, Dianne Gerrelli, Veronica A. Kinsler, Heather C. Etchevers

**Author notes:** Authors contributed to this work equally (Veronica A. Kinsler, Heather C. Etchevers). Current address: University of Southern California, Los Angeles, United States (Magalie E. Carey).

## Abstract

**Background:** MC1R, a G-protein coupled receptor with high affinity for alpha-melanocyte stimulating hormone (αMSH), modulates pigment production in melanocytes from many species and is associated with human melanoma risk. *MC1R* mutations affecting human skin and hair color also have pleiotropic effects on the immune response and analgesia. Variants affecting human pigmentation *in utero* alter the congenital phenotype of both oculocutaneous albinism and congenital melanocytic naevi, and have a possible effect on birthweight.

**Methods and Results:** By *in situ* hybridization, RT-PCR and immunohistochemistry, we show that *MC1R* is widely expressed during human, chick and mouse embryonic and fetal stages in many somatic tissues, particularly in the musculoskeletal and nervous systems, and conserved across evolution in these three amniotes. Its dynamic pattern differs from that of *TUBB3*, a gene overlapping the same locus in humans and encoding class III β-tubulin. The αMSH peptide and the transcript for its precursor, pro-opiomelanocortin (*POMC*), are similarly present in numerous extra-cutaneous tissues. *MC1R* genotyping of variants p.(V60M) and p.(R151C) was undertaken for 867 healthy children from the Avon Longitudinal Study of Parent and Children (ALSPAC) cohort, and birthweight modelled using multiple logistic regression analysis. A significant positive association initially found between R151C and birth weight, independent of known birth weight modifiers, was not reproduced when combined with data from an independent genome-wide association study of 6,459 additional members of the same cohort.

**Conclusions:** These data clearly show a new and hitherto unsuspected role for MC1R in non-cutaneous solid tissues before birth.

## Introduction

The melanocortin-1-receptor (*MC1R*) gene, on human chromosome 16q24.3, encodes a Gs protein-coupled receptor that plays a crucial role in pigmentary phenotype throughout the vertebrate subphylum (Mountjoy et al. 1992; Robbins et al. 1993). It is one of a family of five highly-conserved membrane proteins mediating hormonal responses to the hypothalamic*–* pituitary–adrenal axis throughout the body. On binding, MC1R transduces signals into melanocytes, dendritic cells with specialized organelles called melanosomes, to augment their production of the dark brown eumelanin that colors skin, hair and eyes. Human melanocytes are distributed sparsely but regularly throughout the basal epidermis and hair follicles, from where they transfer their melanosome-bound melanins to designated keratinocytes (Weiner et al. 2007). Melanocyte precursors, derived from highly multipotent and migratory embryonic neural crest cells in all vertebrates (Teillet and Le Douarin 1970), colonize and also synthesize pigment in non-cutaneous sites during development such as the iris, inner ear, heart and central nervous system meninges. In the absence of light or melanosome acceptors, their functions are currently not well understood (eg. Goldgeier et al., 1984; Plonka et al., 2009).

The principal high-affinity ligand of MC1R is α-melanocyte-stimulating hormone (αMSH), a peptide produced from post-translational cleavage of pro-opiomelanocortin (POMC), which is secreted by the anterior pituitary gland but processed locally. Adrenocorticotropin (ACTH) is another POMC cleavage product that can less efficiently activate MC1R, although its principal target is adrenal MC2R (Abdel-Malek et al. 2000; Wikberg et al. 2000). αMSH likewise binds other, lower affinity melanocortin receptors in the central nervous system (Wikberg et al. 2000).

In competent melanocytes, αMSH binding to MC1R triggers a canonical cAMP signal transduction cascade (Newton et al. 2005) that favors enzymatic activity within melanosomes, leading to synthesis of eumelanin rather than the yellow-red pigment pheomelanin from their common metabolic precursor (Hearing and Tsukamoto 1991). This stimulation can be antagonized by members of the agouti signaling protein family, encoded by the single *ASIP* gene in humans, which is conserved in both sequence and function across vertebrates and promotes both reduction of eumelanin synthesis and concomitant pheomelanin production in melanocytes (Suzuki et al. 1997; Yoshihara et al. 2012; Agulleiro et al. 2014).

Partial loss-of-function mutations in *MC1R* have been well tolerated during evolution; modulating its signaling activity leads to a broad pigmentary palette in all vertebrate classes examined (Andersson 2003; Mundy 2005; Gross et al. 2009). In humans, both common (e.g., R151C, R160W, D294H, V92M) and rarer (D84E, R142H) protein alterations from missense variants are associated with reduced eumelanin and proportionately increased pheomelanin synthesis, which leads to lighter skin and red to blonde hair phenotypes (Koppula et al. 1997; Healy et al. 2001). However, many of these variants also confer an increased risk of the pigment cell cancer, melanoma (Valverde et al. 1996; Palmer et al. 2000). The link to oncogenesis is thought to occur in part through variant and isoform-dependent efficacy in stimulating the production and distribution of protective eumelanin, both basally and in response to UV light (Abdel-Malek et al. 2000), while pheomelanin itself also increases oxidative stress in melanocytes (Mitra et al. 2012).

Besides the cyclic AMP signaling pathway, MC1R ligand binding also leads to cross-activation of the MAP kinase family of effectors in both mouse and human melanocytes and in human melanoma. Numerous *MC1R* variants that affect cAMP signaling and eumelanin synthesis do not, however, change the capacity of the proteins they encode to activate the well-characterized effectors ERK1 (MAPK3) and ERK2 (MAPK1; Herraiz et al. 2011). Integration of multiple intracellular effector signals with that of MC1R may therefore affect other aspects of melanocyte physiology than the UV damage-induced tanning response. Indeed, vertebrate integuments are already pigmented at birth, before exposure to UV radiation. *MC1R* variants modify the congenital phenotype of the rare genetic disorders oculocutaneous albinism type 2 (King et al. 2003) and large congenital melanocytic nevus (CMN; Kinsler et al. 2012), strongly suggesting a role for MC1R *in utero* at least in the presence of other mutations affecting pigment synthesis or MAP kinase signaling.

The possibility of non-pigmentary effects *in utero* was raised by an apparent association found between the *MC1R* V92M and R151C alleles and birth weight in both CMN patient and control cohorts (Kinsler et al. 2012). Neural crest cells, the embryonic progenitors of melanocytes, play a key if uncharacterized role in the induction and development of the pituitary (Etchevers et al. 1999; Ueharu et al. 2017), where both POMC and growth hormone are produced. These observations led us to examine how *MC1R* may be a modifier gene for not only rare but common traits.

We therefore explored the hypothesis that MC1R displays pleiotropy before as well as after birth, by examining its expression profile at different times of gestation and in multiple orders of amniotes (bird, rodent, primate). We also compared these findings to the expression pattern of human *TUBB3*, because the first exon of *TUBB3* has been annotated to overlap the coding sequence of *MC1R*, hybrid transcriptional isoforms have been discovered in the pigment cell lineage (Dalziel et al. 2011), and *MC1R* variants could potentially have an effect via either protein product. and *MC1R* variants could potentially have an effect via either protein product. Ultimately, though, we confirmed that human *TUBB3* expression is restricted to the central nervous system before birth. Our data supports an evolutionarily conserved role for the MC1R signaling axis in the development of unexpected tissues like muscle, cartilage, and numerous internal organs.

## Methods

### *In situ* hybridization

Preparation and *in situ* hybridization of human embryonic and fetal material was performed by the Medical Research Council Wellcome Trust Human Developmental Biology Resource with full ethical approval from the National Research Ethics Service, as previously described (Sajedi et al. 2008). Mouse embryos the morning after presumed fertilization were considered to be at embryonic day (E) 0.5, and chick embryos were staged (Hamburger and Hamilton 1992). These were prepared similarly to the human embryos, with fixation in 4% buffered paraformaldehyde in PBS at pH 7.5, paraffin embedding, probe hybridization on 7 µm microtome sections and signal development according to standard protocols (Moorman et al. 2001).

A 539 bp sequence within the 5’UTR of the human *MC1R* mRNA transcript (RefSeq: NM_002386.3) was produced by PCR using F: 5’-GAGCGACGAGATGACTGGAG-3’ and R: 5’-CACAGTCTGTCCTGGTCACC-3’. The PCR product was cloned into the pGEM^®-^T Easy vector (Promega) and digoxigenin-incorporated riboprobes were produced. Sense strand transcription and hybridization was performed on parallel control sections to account for non-specific hybridization signal.

A 439 bp sequence within the *Mus musculus Mc1r* coding region (RefSeq: NM_008559.2;) (Hirobe et al., 2004) was produced by PCR on a genomic DNA template with F: 5’-GACCGCTACATCTCCATCTTCT-3’ and the reverse primer with an additional T7 RNA polymerase recognition sequence (taatacgactcactatagggaga) added at the 5’ end, R: 5’-(T7)AGGAGGAGGAAGAGGTTGAAGT-3’. A 562 bp sequence within the third exon and 3’ UTR of murine *Pomc* (RefSeq: NM_001278582) was similarly amplified with F: 5’-TGACTGAAAACCCCCGGAAG-3’ and R: 5’-(T7)CTAGAGGTCATCAGCTCGCC-3’. A 318 bp sequence within the second exon of *Gallus gallus Mc1r* was amplified with F: 5’-AGCCGACTCCTCGTCCACCC-3’ and 5’-(T7)CACAGCACCACCTCCCGCAG-3’ while a 313 bp fragment of chicken *Pomc* from exon 3 was amplified with F: 5’-GTACCCGGGCAATGGGCACC-3’ and R: 5’-(T7)AGCCGACTCCTCGTCCACCC-3’. *Sox10* probe was synthesized from a pBluescript vector after linearization with HindIII, purification and T3 DNA polymerase as described (Cheng et al. 2000). Purification and *in vitro* transcription of purified T7-extended PCR products were performed as described (Sanlaville et al. 2006). Microphotography was undertaken with an Axioplan2 (human) or Axiozoom (animal) imaging system linked to Zen software (Zeiss). Sequence analysis was checked to make sure of no cross reactivity to other melanocortin receptors.

### Immunohistochemistry

Human embryonic and fetal sections were boiled with Declere^TM^ (Cell Marque) to deparaffinize and rehydrate the tissue, and unmask antigens. Slides were cooled to room temperature and washed using Tris-buffered saline with 0.1% Triton (TBST). Endogenous peroxidase activity within the tissue was quenched using 3% hydrogen peroxidase. Sections were then blocked with 10% calf serum in TBST and incubated with goat anti-MC1R (N-19) polyclonal antibody (1:100 dilution in TBST; sc-6875 Santa Cruz Biotechnology). Sections were then washed and further incubated with a biotinylated secondary polyclonal rabbit anti-goat (IgG) antibody (1:500 dilution in block solution; Dako). Biotin detection was performed using the Vectastain^®^ ABC kit (Vector Laboratories) and diaminobenzidine (DAB) staining. Slides were mounted with VectaMount.

Mouse and chick embryonic and fetal sections were deparaffinized in xylene and rehydrated in phosphate-buffered saline with 0.1% Tween-20 (PBT) with no further antigen unmasking. Endogenous peroxidase activity was quenched as above. Sections were blocked with 2% calf serum in PBT and incubated with rabbit anti-αMSH polyclonal antibody (1:100 dilution in blocking solution; NBT1-78335 Novus Biologicals). After rinsing, a secondary goat anti-rabbit antibody directly conjugated to horseradish peroxidase (111-035-144 Jackson ImmunoResearch) at 1:200 was applied and rinsed after 1h. After DAB staining, slides were counterstained with hematoxylin-eosin before mounting with Eukitt.

### Reverse Transcriptase-PCR

Human microdissected tissues at defined stages were homogenized and whole RNA was extracted using the Trizol method according to manufacturer’s recommendations (Thermofisher). cDNA was synthesized using the M-MLV reverse transcriptase kit (Promega) and assessed for the absence of genomic DNA contamination. PCR was carried out using standard conditions using cDNA with the following gene specific primers: *MC1R-*Fwd-ACTTCTCACCAGCAGTCGTG, *MC1R-*Rev-CATTGGAGCAGACGGAGTGT. *TUBB3-*Fwd-TCTACGACATCTGCTTCCGC and *TUBB3-*Rev-TCGTCTTCGTACATCTCGCC.

### Genotyping Analysis

#### Samples

867 phenotyped ALSPAC (Avon Longitudinal Study of Parent and Children http://www.bristol.ac.uk/alspac/) DNA samples from “Northern European/Caucasian/white” children were assessed. Ethical approval for the study was obtained from the ALSPAC Ethics and Law Committee and the Local Research Ethics Committees. The entire *MC1R* gene was sequenced using standard Sanger sequencing and Big Dye Terminator v3.1 under manufacturers’ specifications (ThermoFisher, UK). Genotyping was carried out by sequence trace analysis using Sequencher software (Softpedia). The additional genotyping data for R151C was obtained from ALSPAC upon request.

#### Statistical Analysis

Logistic regression modeling was used to test the hypothesis that MC1R genotypes R151C or V92M were statistically associated with increased birth weight in the population samples examined. The covariates added to the model were sex, gestation (in completed weeks), maternal pre-pregnancy weight (kg) and smoking (tobacco smoked in the last two weeks). Only babies born at term (between 37 and 42 weeks) were used in this study. Analysis was performed using the SPSS statistical package (version 21).

## Results

### Expression of MC1R during human embryonic and early fetal development

We analyzed MC1R expression in normal human tissues from embryos and fetuses at Carnegie stages (CS) 18 (6-7 weeks’ gestation [wg]), CS23 (8 wg), Fetal stage 1 (F1 - early 9 wg), F3 (11 wg) and 18 wg.

*In situ* hybridization revealed widespread transcription of *MC1R* mRNA in the skeletal system, demonstrated in the cells, periosteum, and muscle fibers surrounding the femur and patella, and the forelimb radius and ulna, at CS23 (**Figure 1A**). In addition, *MC1R* transcripts were observed in the liver, pancreas, adrenal cortex, and the tubules and glomeruli of the renal cortex at CS23, as well as the bronchial epithelia of the lung and semicircular canals of the inner ear (**Figure 1B; Supplemental Figure 1**). *MC1R* mRNA was likewise detected throughout the pituitary, particularly the anterior lobe, in the ventricular zone lining the forebrain ventricles (**Figure 1C**), and in the neural retina and ganglionic eminences (**Supplemental Figure 1**).

**Figure 1.**
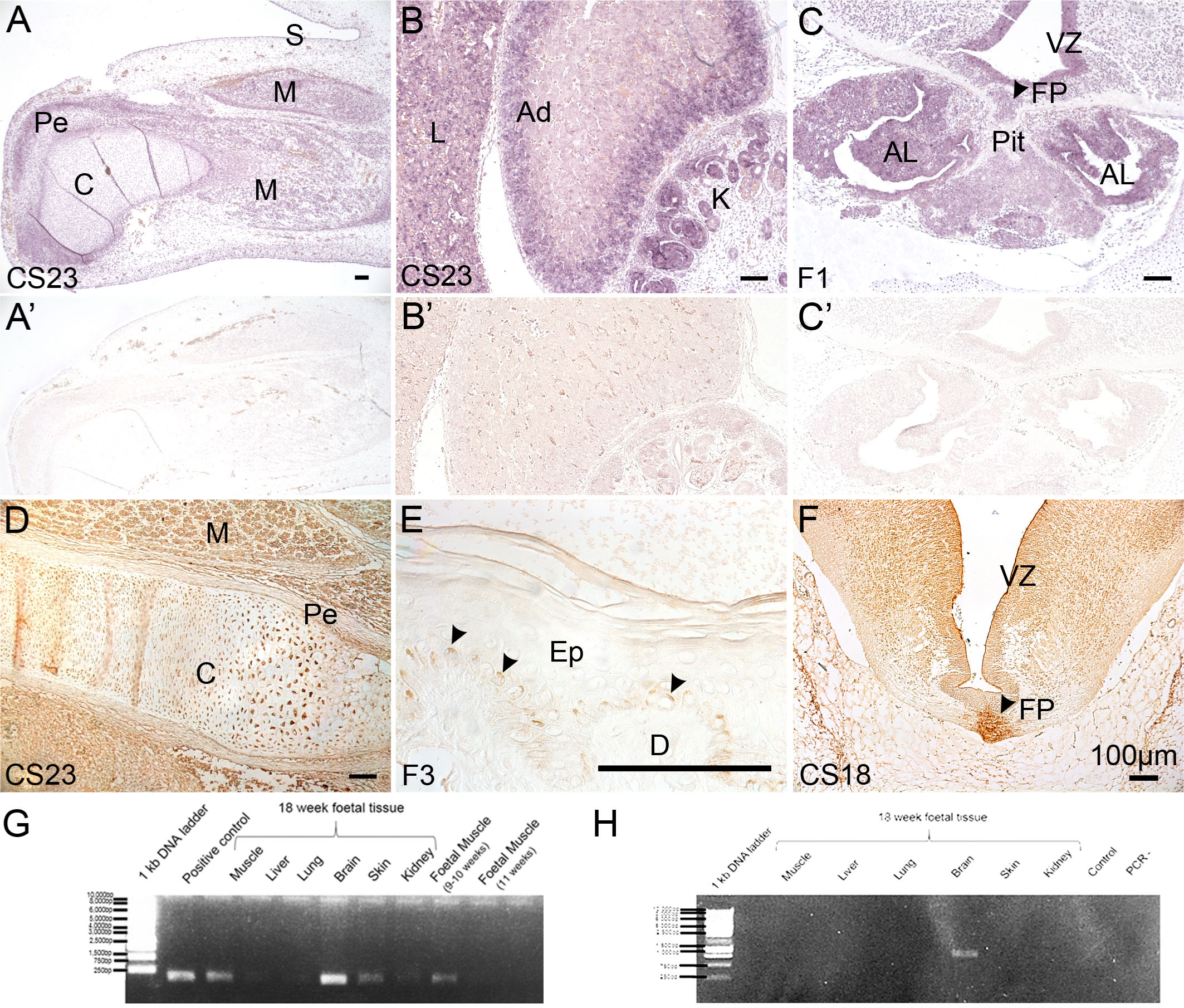
Human MC1R expression in staged late embryonic (CS18, CS23) and early fetal tissues (F1, F3). **A-C**, *in situ* hybridization; **A’-C’**, hybridization of adjacent sections to sense-transcribed probe (negative controls). D-F, immunohistochemistry without counterstain. **A**. Humerus cartilage, particularly its perichondrium, skeletal muscle and skin all transcribe *MC1R* at CS23. B. *MC1R* transcripts were observed in the liver, the adrenal cortex, and the glomeruli of the renal cortex at CS23. **C**. Coronal section through the ventral diencephalon at fetal stage 1 (end of 8 weeks’ gestation) demonstrating *MC1R* mRNA in the ventricular zone and floorplate, the parenchyme, and throughout the pituitary gland, with strong expression in the anterior lobe. **D**. Protein expression in the musculoskeletal system, including the perichondrium and cartilage of the femur and its surrounding muscles at CS23. **E**. Fetal skin at stage 3, approximately 10 weeks’ gestation, shows restricted MC1R protein (arrowheads) on the apical aspect of cells in the proliferative layer of the basal epidermis. No expression is observed in the dermis or in upper epidermis, although non-specific signal is trapped in the outermost corneal layer. **F**. At CS18 (approximately 7 weeks’ gestation), MC1R protein is already present in neuron cell body tracts and nuclei of the hindbrain, in the ventricular zone, and particularly concentrated in the floorplate at this level. **G**. RT-PCR of *MC1R* cDNA in fetal tissues at 18 wg, F1 (9 wg) and F3 (11 wg). **H**. RT-PCR of *TUBB3* cDNA in 18 wg tissues. Ad, adrenal gland; AL, anterior lobe; C, cartilage; CS, Carnegie stage; D, dermis; F, fetal stage; Ep, epidermis; FP, floorplate; K, kidney; L, liver; M, muscle; Pe, perichondrium; Pit, pituitary; S, skin; VZ, ventricular (proliferative) zone of central nervous system. Bars = 100 µm.

Using immunohistochemistry, we confirmed expression of MC1R protein in the chondrocytes and periosteum of the developing femur at CS23, as well as in surrounding skeletal muscle (**Figure 1D**). Positive staining was also seen in the epiphysis of the CS23 humerus (not shown). MC1R was distributed in the basal layer of the epidermis (**Figure 1E**) at F3, preceded by expression in the CS18 floor plate and ventricular zone of the brain (**Figure 1F**).

We then validated the presence of *MC1R* expression by RT-PCR in cDNA derived from later fetal tissues. Expression was detected in a F3 (9-10 wg) muscle sample, although not in one from 11 wg. At 18 wg, muscle, brain, skin and kidney all transcribed *MC1R*, in contrast to samples from liver and lung (**Figure 1G**).

During this study, it became important to be able to distinguish between human *MC1R* and *TUBB3* transcription. *TUBB3* is known to be primarily expressed in neurons and fetal astrocytes, where it is involved in microtubule formation (Dráberová et al. 2008). Due to a “leaky” polyadenylation site between *MC1R* and its 3’ neighbor *TUBB3*, it has been shown that transcripts containing the whole of *MC1R* fused to *TUBB3* can also be expressed in human, although not mouse, melanocytes (Dalziel et al. 2011; Herraiz et al. 2015). Two distinct antibodies can recognize either endogenous or chimeric TUBB3 protein in human melanoblasts and melanocytes *in vitro* and *in vivo* (Locher et al. 2013). For these reasons, we confirmed that *in situ* hybridization reflected endogenous human *MC1R* expression by validating with immunohistochemistry and RT-PCR. As expected from the known function of β3-tubulin, *TUBB3* mRNA expression was only detected in the 18 wg brain sample (**Figure 1H**).

### Analysis of *Mc1r* expression in mouse embryonic and fetal development

To compare developmental expression of MC1R with that of other species in which additional fetal stages and tissues could also be examined, we performed *in situ* hybridization of antisense *Mc1r* probe to mouse sections from embryonic day (E)11.5, E13.5, E17.5, E19.5, postnatal day 2 and adult hairy skin. Mc1r protein expression was also examined by immunohistochemistry at E13.5.

At E11.5, the dorsal root and cranial ganglia and the ventricular zone of the central nervous system (CNS) already expressed Mc1r. It was also transcribed at relatively lower signal intensity throughout early skeletal muscle and mesonephros and many other forming tissues (**Figure 2A, B**). By E13.5, expression appeared strong within liver, spinal cord and dorsal root ganglia, and limb perichondrium and muscle (**Figure 2C**). The hindbrain, pituitary gland and future cochlea transcribed *Mc1r*, in addition to tongue, neck and vascular smooth muscle (**Figure 2D**). Mc1r protein at this stage was present in the heart and arterial smooth muscle, esophageal and lung segmental bronchial epithelia (but not in main bronchi), and concentrated in the CNS at the floorplate and in grey matter (**Figure 2E, F**). In contrast to widespread transcription in limb, head, rib and pelvic cartilage, Mc1r was detectable in the intervertebral discs (Figure 2E) but not in the vertebral body itself (**Figure 2E, F**).

**Figure 2.**
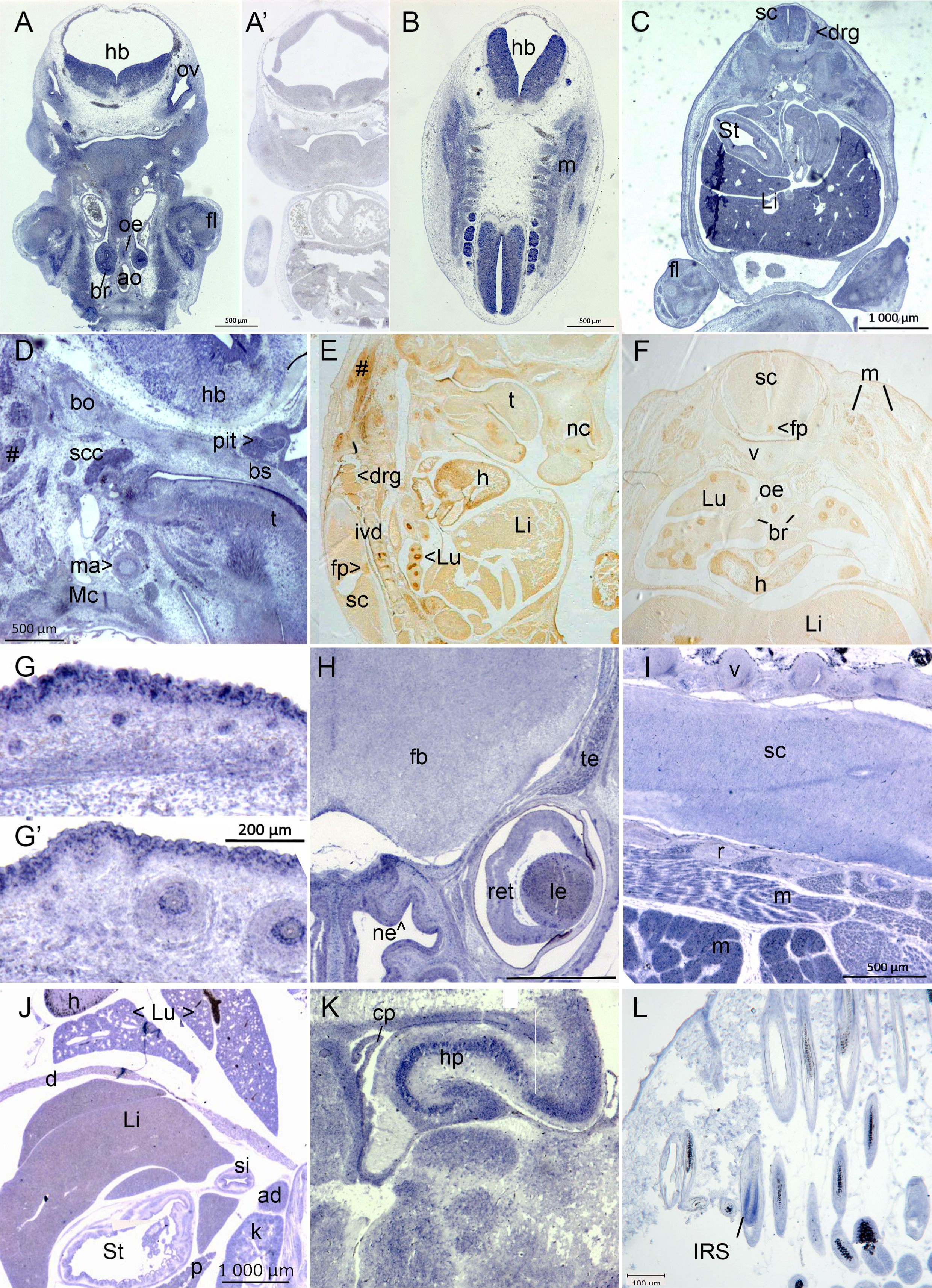
Embryonic mouse expression of *Mc1r* RNA (blue) and Mc1r protein expression (brown). **A, B.** Embryonic day of gestation E9.5, frontal sections, antisense probe. **A’**. Hybridization of adjacent section to sense-transcribed probe (negative control). **C**. Mc1r expression is widespread at E13.5. **D**. Facial, head and neck tissues at E13.5. E, F. Immunohistochemistry in sagittal and transverse section respectively at E15.5. **G, G’**. Hair follicles of skin (**G**) and whisker vibrissae (**G**’) at E17.5. **H**. Frontal section of head, E17.5. **I.** Oblique section through dorsal trunk, E17.5. **J**. Frontal section through ventral trunk, E17.5. **K**: Postnatal day 2 forebrain. **L**. Adult skin, hair follicles with black pigment sheath. Abbreviations: ad, adrenal gland; ao, aorta; br, bronchi; bo, basioccipital cartilage; bs, basisphenoid; cp, choroid plexus; d, diaphragm; drg, dorsal root ganglion; fb, forebrain; fl, forelimb; fp, foreplate; h, heart; hb, hindbrain; hp, hippocampus; IRS, inner root sheath; ivd, intervertebral disc; k, kidney; Li, liver; le, lens; Lu, lung; m, muscle; ma, mandibular artery; Mc, Meckel’s cartilage; nc, nasal cartilage; ne, nasal epithelium; oe, oesophagus; ov, otic vesicle; p, pancreas; pit, pituitary; r, rib; ret, retina; sc, spinal cord; scc, semi-circular canal; si, small intestine; St, stomach; t, tongue; te, temporalis muscle; v, vertebra.

Additional epithelia transcribing *Mc1r* in late gestation included guard hair follicles and epidermis (Hirobe et al. 2004)(**Figure 2G**), vibrissae (**Figure 2G’**), and the lining of the nasal cavity at E17.5 (**Figure 2H**). The nasal cartilage and frontal bones also expressed *Mc1r* as well as the temporalis, orbicularis oculi (**Figure 2H**) and zygomatic facial muscles. Expression remained strong in the neurosensitive retina as well as in CNS neurons throughout the brain, including the forebrain and spinal cord (**Figure 2H, I**). The trapezius, hindlimb muscle groups, and intercostal muscles transcribed *Mc1r* (**Figure 2I**).

Smooth muscle layers of the stomach (E13.5) and diaphragm (E17.5) were positive (Figure 2C, 2H), and expression also continued in the liver, lung and heart (**Figure 2J**) as well as the thymus (not shown). By postnatal day (P)2, strong transcription of *Mc1r* was observed in pyramidal neurons of the hippocampus and at lower levels in motor nuclei, a subependymal layer of the cerebral cortex, and the choroid plexus (**Figure 2K**). The skeletal muscle of the hypodermis (*panniculus carnosus*, not shown) as well as follicular keratinocytes of the inner root sheath, but not the interfollicular epidermis, express *Mc1r* in adult skin (**Figure 2L**).

### Analysis of *Mc1r* expression in chicken embryonic and fetal development

*Mc1r* was not detected during the first few days of development, at stages (HH; Hamburger and Hamilton 1992) 11, 16 and 17 (not shown). By HH24-25, weak expression appeared throughout the neural tube (brain and spinal cord), somitic myotome and mesonephros (**Figure 3A**) as well as in cranial ganglia, limb and pharyngeal mesenchyme (**Figure 3B**). More intense zones of transcription appeared in subectodermal lateral plate or limb mesoderm (arrowheads, **Figure 3A, B**), not corresponding to specific anatomical features. *Mc1r* was more widely transcribed by early fetal stage HH29, corresponding to 6-6.5 days’ incubation. Sites included the ectoderm and intercostal and epaxial skeletal muscles (**Figure 3C**), the salivary gland, and some cephalic muscles including oculomotor and pterygoid. Expression was observed in the brain and cranial ganglia with comparatively strong expression in discrete zones of subectodermal mesenchyme of both the beak primordium (**Figure 3D**, arrows) and forelimb (**Figure 3E**, arrow), but little to none in the cartilage itself.

**Figure 3.**
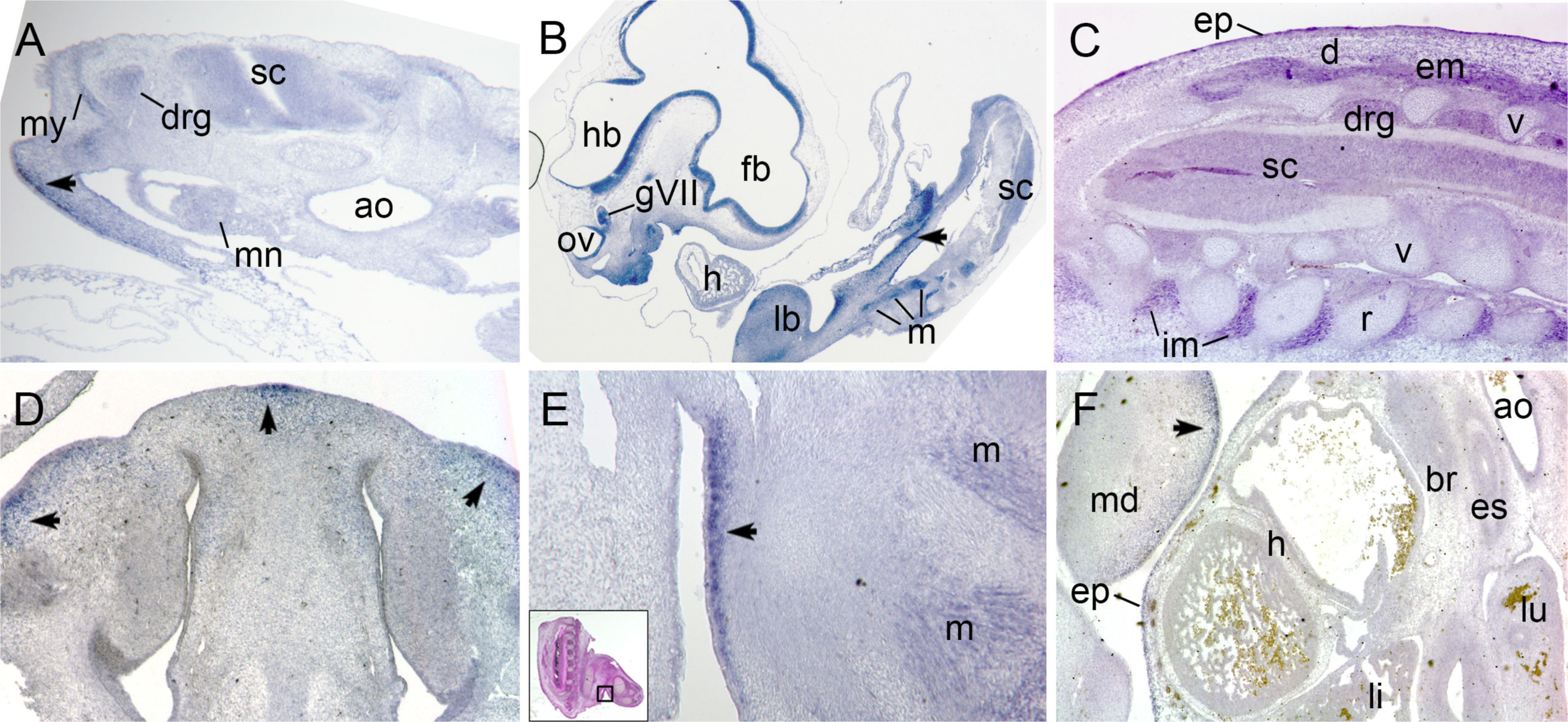
Chicken *Mc1r* RNA transcription (blue) during embryonic stages. **A**. Oblique transverse section through trunk at HH24. **B**. Parasagittal section through whole embryo, HH25. **C**. Oblique transverse section through dorsal trunk, HH29. **D**. Coronal section through nasal cartilage, presumptive upper beak, HH29. **E**. Ventral forelimb, area magnified indicated on inset (hematoxylin-eosin stain), HH29. **F**. Parasagittal section through upper chest cavity, HH32. Abbreviations: ao, aorta; br, bronchus; d, dermis; drg, dorsal root ganglia; em, epaxial muscle; ep, epidermis; es, esophagus; fb, forebrain; h, heart; hb, hindbrain; gVII, acoustic ganglion; im, intercostal muscle; lb, forelimb bud; li, liver; lu, lung; m, muscle; md, mandible; mn, mesonephros; my, myotome; ov, otic vesicle; r, rib; sc, spinal cord; v, vertebra.

By HH32 (approximately 7.5-8 days’ incubation), skeletal muscles and localized perichondrial *Mc1r* expression continued in both limb and trunk (**Figure 3F**). Areas of mesenchyme also continued to transcribe *Mc1r*, in which case the ectoderm expressed comparatively less transcript (arrow); tracheal smooth muscle had some, but kidney (not shown), liver and heart showed little to no *Mc1r* expression at this stage.

### Ligand expression

*POMC* transcripts are translated into the peptide pro-opiomelanocortin, which is enzymatically processed to yield two hormones that are each agonists of MC1R (Suzuki et al., 1996): the higher affinity alpha-melanocyte-stimulating hormone (α-MSH), and adrenocorticotropic hormone (ACTH). In order to establish whether Mc1r is likely to be activated before birth by specific ligand-dependent binding, we examined the transcription of *Pomc* during mouse and chicken embryogenesis by *in situ* hybridization.

An antisense RNA probe against murine *Pomc* at E11.5 showed widespread expression throughout the CNS, in developing skeletal muscle masses, cartilage and craniofacial mesenchyme (**Figure 4A**). Stronger localized expression was seen by E13.5 in the dorsal root ganglia, spinal cord and lung (**Figure 4B**) as well as liver, dermis, epaxial skeletal muscle but also diaphragmatic and intestinal smooth muscle, but less intensely in cartilages such as the vertebral body (**Figure 4C**). In the head at the same stage (**Figure 4D**), *Pomc* remained widely transcribed, as in the heart (cf. the mouse heart’s *Mc1r* expression in Figure 2E, J). Epithelia in close contact with overlying mesenchyme such as the salivary glands, the nasal epithelia, the stomach and the lung were all positive (**Figure 4C, D**). *Pomc* also showed intense expression in areas of the tongue, nasal epithelium, pons, cerebellum and hypothalamus. A complementary coronal section through a chicken embryo at HH31 also showed high levels of transcription within a subset of hypothalamic neurons and transcription through the hindbrain (**Figure 4E**). The nasal glands and epithelium, facial muscles and vibrissae, as well as more standard-sized hair follicles of the face (**Figure 4F**) expressed *Pomc* at E17.5 in the mouse. Within the central nervous system, the neural retina, to a much lesser extent the glia of the optic nerve, the hippocampus and cerebellum continued to strongly transcribe *Pomc*.

**Figure 4.**
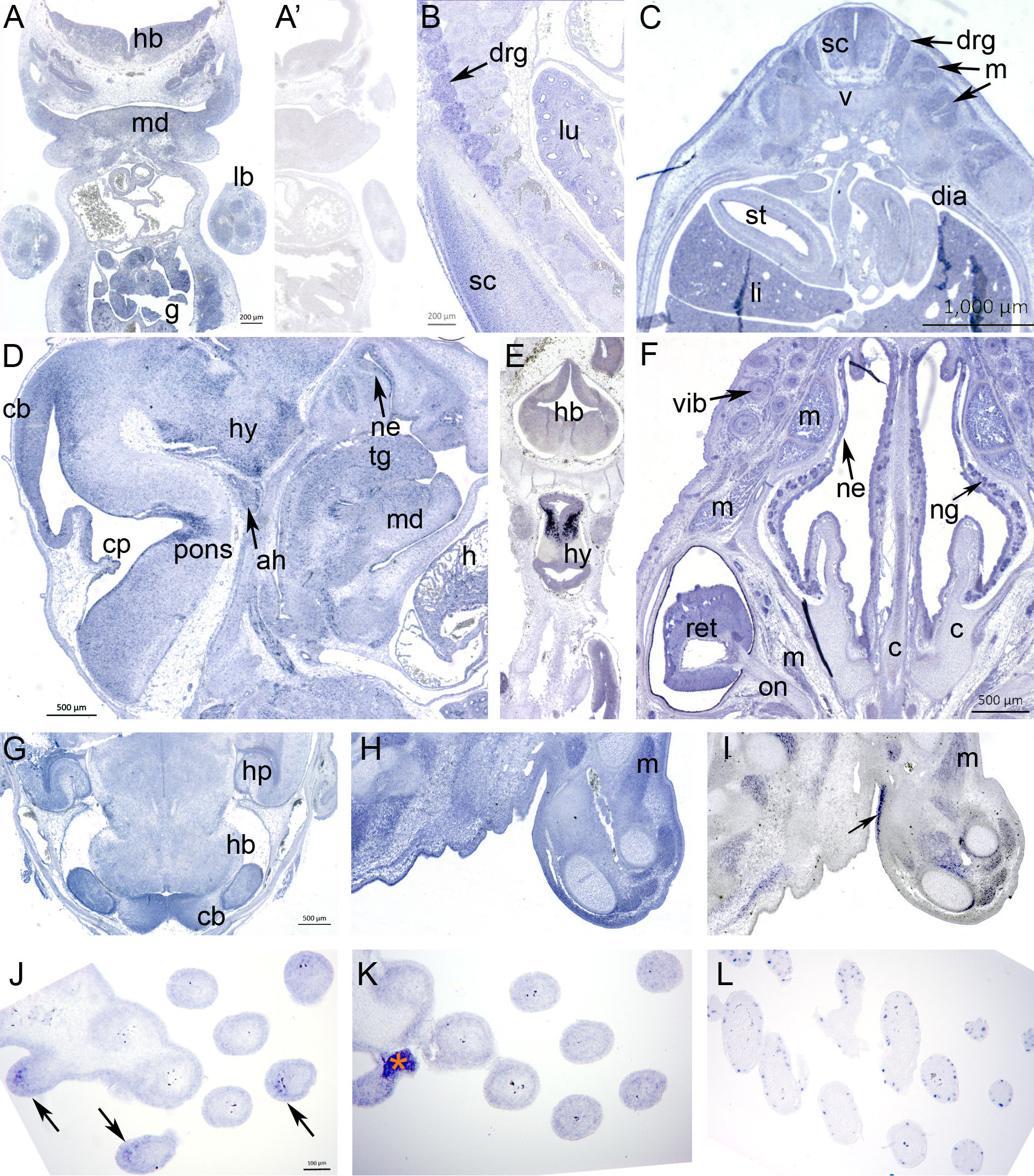
Embryonic mouse *Pomc* transcription and embryonic chicken *Pomc* expression as relates to *Mc1r* and/or *Sox10* transcription (blue). **A**. Frontal section, E11.5; strong expression in rib cartilage and intercostal and limb muscle masses, mandibular mesenchyme, and gut. **A’**. Hybridization of adjacent section to sense-transcribed probe (negative control). **B**. Oblique parasagittal section through upper dorsal trunk at E13.5; strong expression in central and peripheral nervous systems and lung mesenchyme. **C**. Transverse section through trunk at same stage, showing widespread additional transcription by smooth and skeletal muscle and liver. **D**. E13.5 head in sagittal section, showing hypothalamic, cerebellar, pontine and pituitary transcription of *Pomc*, as well as in heart and tongue muscle. **E**. Coronal section of an embryonic chicken at stage HH31 with extremely strong transcription by hypothalamic neurons in addition to expression throughout the brain. **F**. Mouse E17.5 coronal section through the level of the eye with striking *Pomc* expression in the neural retina, facial muscles but not cartilage, and nasal glands. **G**. The hypothalamus, cerebellum and select hindbrain nuclei express relatively more *Pomc* transcript at E17.5. **H**, **I**: Adjacent sections of chicken forelimb at HH29, hybridized respectively with probes against Pomc and Mc1r. Pomc (**H**) is more broadly expressed throughout connective tissues, cartilage and the dermis and intense staining is present in skeletal muscle. **I**. *Mc1r* is also expressed in muscle masses but also regions of perichondrium and a specific zone of dermis (arrow) not anatomically distinguished. **J-L**: Tangential sections through wing skin and feather follicles at HH37, with probes against (**J**) *Pomc* - asymmetric dermal expression indicated by arrows, (**K**) *Mc1r* - artefact labelled with orange asterisk - or (**L**) *Sox10* (melanoblasts). Abbreviations: ah, adenohypophysis; c, cartilage; cb, cerebellum; cp, choroid plexus; dia, diaphragm; drg, dorsal root ganglon; g, gut; h, heart; hb, hindbrain; hp, hippocampus; hy, hypothalamus; lb, forelimb bud; li, liver; lu, lung; m, muscle; md, mandible; ne, nasal epithelium; ng, nasal gland; on, optic nerve; pons, pontine flexure; ret, retina; sc, spinal cord; st, stomach; tg, tongue; v, vertebra; vib, vibrissa.

We compared the expression domains of *Pomc* to *Mc1r* in the HH29 chicken forelimb (**Figure 4H, I**; also *cf.* Figure 3E). While Pomc was widely expressed, Mc1r was also widely expressed, with lesser transcription in connective tissue throwing muscle and perichondrial transcription into higher relief. Strong expression in a region of junctional dermis on the proximal ventral face for *Mc1r* (**Figure 4I**, arrow) was complementary to strong dermal transcription of *Pomc* under the rest of the limb epidermis (**Figure 4H**). Both *Pomc* and Mc1r continued to be expressed in feather follicular epidermis and dermis, with an asymmetric distribution of *Pomc* in the same dermal compartments seen in cross-section (**Figure 4J, K**). *Mc1r* also appears to be in feather melanoblasts at this age, highlighted by their specific transcription of the transcription factor *Sox10* (**Figure 4L**), at the onset of the period when the chicken fetus begins to show pigmentation of some maturing follicles.

Immunohistochemistry against αMSH (**Figure 5**) showed that most but not all embryonic sites of *Pomc* transcription yielded the presence of this specific ligand in chicken or mouse. At murine E13.5, αMSH was synthesized in limb epidermis, muscle masses, perichondrium and cardiomyocytes (**Figure 5A-B**). By E17.5, immunoreactivity remained strong in limb skin and skeletal muscle (including the *panniculus carnosus*) but not in the cartilage or perichondrium (**Figure 5C**). The hormone was also produced by the E17.5 liver, pancreas, intestinal epithelium, adrenal gland and kidney (**Figure 5D, E**), as well as interdigital mesenchyme (**Figure 5F**). In the chicken, strong immunoreactivity was observed in the ocular choroid plexus, between the retinal pigmented epithelium and the (negative) sclera (**Figure 5G**), at HH29. As in the mouse limb and paw, synthesis was excluded from embryonic chicken cartilage at this stage in both proximal limb and wingtip, as well as from the dorsal root ganglia. However, αMSH immunoreactivity was observed in perineural sheaths and distal nerves; subectodermal mesenchyme in a manner reminiscent of, but not superimposing, *Mc1r* transcription in body wall and limb (cf. Figure 3), and to a lesser extent in interdigital mesenchyme (**Figure 5H**).

**Figure 5.**
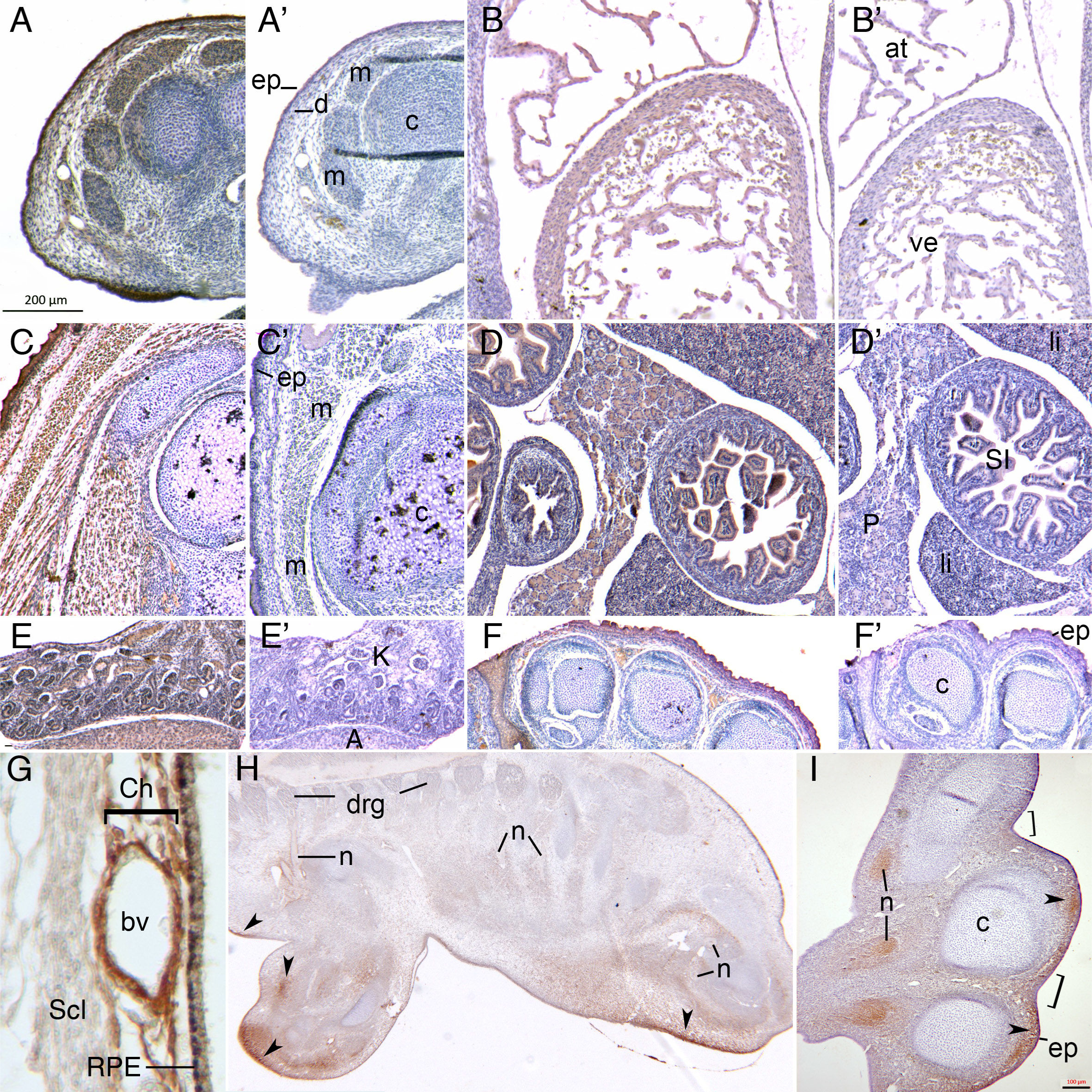
Immunolocalization of alpha melanocyte-stimulating hormone (αMSH) in mouse and chicken embryonic tissues. **A-F**: Each had an adjacent section counterstained but incubated without primary antibody as a negative control (**A’-F’**). **A**. Mouse E13.5 limbs show αMSH in skin, muscles and perichondrium. B. At the same stage, αMSH in the heart. C. By E17.5 in the mouse limb, αMSH production is strong in the epidermis but also in the dermis and all skeletal muscle, but excluded from hypertrophic cartilage. **D, E**. αMSH is unexpectedly produced by numerous internal organs at the same stage. F: In addition to the skin, interdigital mesenchyme before digit separation is complete, expresses αMSH, in contrast to the digital cartilage. G. Chicken embryonic eye at HH29 shows strong choroidal expression of αMSH in close proximity to the blood vessels and retinal pigmented epithelium. H. At the same stage, immunoreactivity was observed throughout the epidermis but also in discrete limb and flank subectodermal dermis regions (arrowheads), in perineural sheaths but not in dorsal root ganglia. **I**. αMSH is produced in distal nerves of the forelimb at HH29, in the epidermis with the exception of interdigital skin (brackets). As in the trunk, αMSH is produced by specific domains of underlying dermal mesenchyme at the tips of the growing digits. Abbreviations: A, adrenal; at, atrium; bv, blood vessel; c, cartilage; Ch, ocular choroid; d, dermis; drg, dorsal root ganglia; ep, epidermis; K, kidney; li, liver; m, muscle; n, nerves; P, pancreas; RPE, retinal pigmented epithelium; Scl, sclera; SI, small intestine; ve, ventricle.

### Birth weight analysis using logistic regression modelling

In an earlier study, we found that of 270 normal children from the English ALSPAC study (Jones et al. 2000), 42 carried at least one R151C variant and 39 at least one V92M variant of *MC1R*. These genotypes were independently significantly associated with increased birth-weight (p = 0.002), and independent of known birthweight modifiers (Kinsler et al. 2012). We therefore genotyped 867 children from the ALSPAC cohort for these two alleles, and confirmed a significant positive association (p = 0.05) between the R151C variant and birth weight in this group, again independent of other known modifiers (**Table 1**). Within this larger sample, *MC1R* R151C-variant newborns had a mean birthweight that was 74 g grams heavier than other genotypes, including wild type and other variants. However, V92M no longer showed a significant association with birth weight (p = 0.41). When combined with GWAS data using 6,459 additional individuals from the same cohort (total samples analysed 7,326), the R151C association also was no longer significant (p = 0.40) (**Table 1**). Thus, variations in *MC1R* genotype and, by extension, signaling activity, do not appear to correlate with birth weight in the general English population as represented by the ALSPAC cohort. A similar analysis was then performed using maternal *MC1R* genotype at these two alleles, and birthweight of the child. We had postulated that a connection between maternal genotype and Vitamin D status could have nutritional effects on growth of the fetus. However this analysis also failed to demonstrate a significant association.

**Table 1:**
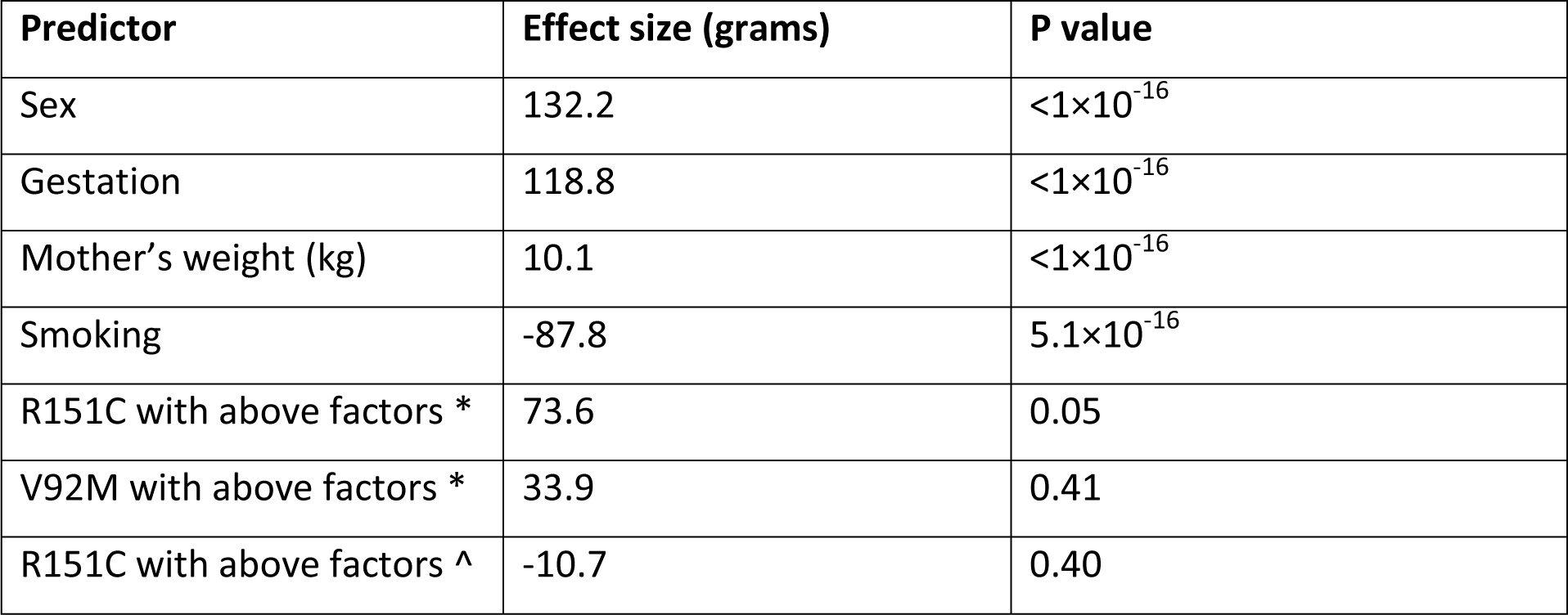
Association of *MC1R* genotype with birth weight. *MC1R* R151C is positively associated with heavier birth weight (p<0.05) using in-house Sanger sequenced genotyping data obtained from 867 subjects, but this association disappears (p=0.40) when combined with ALSPAC GWAS data from an additional 6459 individuals (n=7,326 normal births). V92M has no association with birth weight using in-house genotyping data (GWAS data unavailable for V92M). Covariates known to affect birth weight (infant sex, total gestation in weeks, pre-pregnancy weight in kilograms and level of tobacco smoke) were added to the model. *Data available from 867 initial in-house samples ^Data from 7,326 individuals (867 from in-house genotyping plus 6459 ALSPAC GWAS data)

| Predictor | Effect size (grams) | P value |
| --- | --- | --- |
| Sex | 132.2 | $<1 \times 10^{-16}$ |
| Gestation | 118.8 | $<1 \times 10^{-16}$ |
| Mother's weight (kg) | 10.1 | $<1 \times 10^{-16}$ |
| Smoking | -87.8 | $5.1 \times 10^{-16}$ |
| R151C with above factors * | 73.6 | 0.05 |
| V92M with above factors * | 33.9 | 0.41 |
| R151C with above factors ^ | -10.7 | 0.40 |

## Discussion

Our previous study led us to consider functional roles for MC1R before birth that may affect weight at birth, and thereby to investigate its expression in the developing embryo. In this work, we have demonstrated evolutionary conservation of previously-undescribed *Mc1r* and *MC1R* expression domains, particularly in specific compartments of the prenatal skin and skeletal musculature, in three amniote species including humans. In addition, the widespread transcription of *Pomc* we document for the first time in the developing tissues of both avian and murine embryos, supports an evolutionarily conserved role for Mc1r in regulating growth of multiple organ systems, including those potentially affecting weight at birth. Investigation of potential correlations between two variant *MC1R* genotypes and increased birth weight in human infants did not allow us to further support the hypothesis that these alleles had a measurable effect on this specific phenotype for the general population. Intriguingly, many years ago, strong correlation between light hair and freckling, traits known to be influenced by *MC1R* genotype, and “body build” (an early type of BMI relating height to weight) had already been noted (Brues 1950). Model systems may better lend themselves to future investigations of whether body mass is influenced by Mc1r activity in humans or other animals.

Nonetheless, data presented here clearly show that although amniote embryos are pigmented at birth, the αMSH-Mc1r signaling axis plays roles well beyond the promotion of postnatal melanogenesis in the three species examined here. In humans, pleiotropic effects of MC1R are already known in post-natal life. UV stimulation of αMSH production in the epidermis leads to paracrine MC1R binding and activation in interfollicular melanocytes to promote the synthesis of eumelanin and, ultimately, a tanning response *in vivo*, but also to their population expansion *in vitro* (Valverde et al. 1995; Abdel-Malek et al. 2000). Women with two variant *MC1R* alleles show greater pain relief in response to the κ-opioid pentazocine; Mclr mediates κ-opioid analgesia in mice as well (Mogil et al., 2003). Later work demonstrated overall decreased nociception in both humans and mice bearing functionally variant *MC1R*/*Mc1r* alleles that decrease eumelanin synthesis (so-called “red hair” alleles), implying that endogenous activation of MC1R may counteract µ-opioid-mediated analgesia and confer greater basal pain sensitivity to non-carriers of both sexes (Mogil et al., 2005). MC1R transfected into immortalized human embryonic kidney cells leads to an apparent agonist-independent increase in cAMP levels (Sanchez-Más et al. 2004). We have found that *Pomc* is normally transcribed in the fetal mouse kidney and that human embryonic kidney strongly expresses *MC1R*, supporting a preponderant role for ligand-dependent stimulation *in vivo*, be it autocrine, paracrine or endocrine.

Human primary chondrocytes isolated from osteoarthritic adult knees express αMSH-responsive MC1R, and signal through the cAMP pathway to induce the synthesis of collagens as well as some MMP degradation enzymes (Grässel et al. 2009). A chondrosarcoma cell line also expresses MC1R and reduces levels of post-inflammatory *MMP13* transcription in response to αMSH, while healthy primary chondrocytes do not. Our observation of normal expression of *Mc1r* in the perichondrium in embryonic chick, mouse and human cartilage, and of the protein within fetal chondrocytes, supports the hypothesis that MC1R activity normally promotes growth and matrix remodeling of developing cartilage. More intriguingly yet, we observed strong MC1R receptor expression in the developing human heart, kidney, adrenal gland, liver, pancreas and lung.

In mice, where inter-follicular melanocytes rapidly disappear after birth from most sites and postnatal pigmentation is thereafter restricted to a self-renewing population within hair follicles, Mc1r stimulation by αMSH favors the terminal differentiation of as yet unpigmented melanoblasts (Hirobe 1992). We have now shown that *Mc1r* is widely expressed in the pre- and perinatal epidermis but restricted by P2 to the follicular bulb and sheath within which melanoblasts are embedded, suggesting that both epidermal components develop in synchronized but cell-specific manners in response to hormonal stimulation. In addition, *Mc1r*-null mice are less resistant to experimentally induced oxidative stress and inflammation, leading to dermal fibrosis and collagen synthesis or the aggravation of colitis (Böhm and Stegemann 2014). In humans, the R163Q allele of *MC1R*, which leads to reduced cAMP signaling upon αMSH binding, is also strongly associated with susceptibility to hypertrophic scarring (Sood et al. 2015).

Finally, in the developing chicken, we have shown that feather germ keratinocytes express *Mc1r* by mid-gestation, when *Sox10*+ melanoblasts are present within the sheath and interfollicular epidermis. *Sox10* is a transcription factor that is expressed by multipotent neural crest cells, repressed temporarily upon their colonization of the epidermal annexes, and re-expressed by transiently amplifying and terminally differentiated melanocytes (Osawa et al., 2005). The *Pomc* transcript is expressed asymmetrically in cross-section of the dermal pulp. Interestingly, the Asip antagonist is also asymmetrically translated later, within the postnatal dermal pulp compartment, facing, and likely regulating the color of, forming feather barbs (Yoshihara et al. 2012).

In conclusion, we have identified *MC1R* expression in a range of unreported tissues in the developing human, chick and mouse embryo and fetus. Expression of both ligand and receptor in multiple sites of subepithelial mesenchyme and in many similar tissues and organs suggest that MC1R has widespread unknown functions in fetal growth and differentiation evolutionarily conserved across species.

## Tables and Figure Legends

**Supplemental figure 1.**
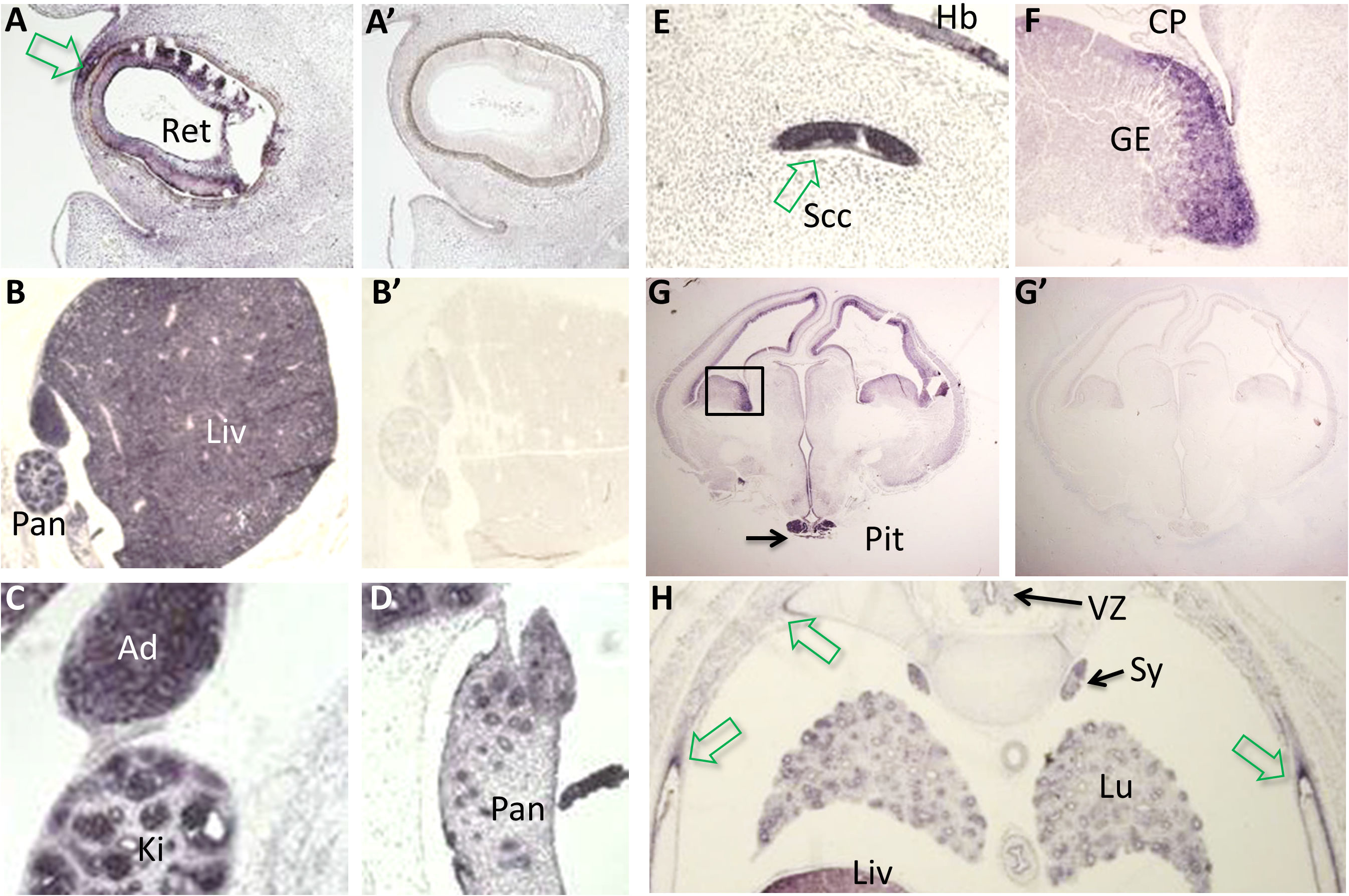
Additional expression domains at late embryonic stages of human *MC1R*. **A, B, G**: Each had an adjacent section hybridized to sense-transcribed probe, as a negative control (**A’, B’, G’**). **A**: The neural retina and corneal mesenchyme (arrow) expresses *MC1R* much more than the overlying corneal epithelium. **B-D**: Liver, kidney and adrenal gland, and pancreas - particularly duct epithelium - all strongly expressed *MC1R*. **E, F**: The semicircular canal of the inner ear, the hindbrain and the forebrain ganglionic eminence were all positive, and to a lesser extent, the choroid plexus. **G**: Region from which **F** was derived in box; the forebrain ventricular zone and the pituitary gland strongly transcribe *MC1R*. **H**: Lung epithelium but also mesenchyme, sympathetic ganglia, the liver and what appear to be tendon attachment points or perichondrium of the ribs (arrowheads) all express *MC1R*. Ad, adrenal gland; CP, choroid plexus; GE, ganglionic eminence; Hb, hindbrain; Ki, kidney; Liv, liver; Lu, lung; Pan, pancreas; Pit, pituitary; Ret, retina; Scc, semicircular canal; Sy, sympathetic ganglia; VZ, ventral zone.

## Acknowledgements

We are extremely grateful to all the families who took part in this study, the midwives for their help in recruiting them, and the whole ALSPAC team, which includes interviewers, computer and laboratory technicians, clerical workers, research scientists, volunteers, managers, receptionists and nurses. We thank Sarah Lechat and Julia Matonti for their technical assistance. Chicken *Sox10* plasmid was graciously provided by Paul Scotting. The UK Medical Research Council and the Wellcome Trust (Grant ref: 092731) and the University of Bristol provide core support for ALSPAC. Caring Matters Now (VK, HCE); Nevus Outreach, Inc. (MC, HCE); the RE(ACT) Community (PH, HCE); Asociación Española de afectados por Nevus Gigante Congénito (HCE); Nævus 2000 France-Europe and the Association du Nævus Géant Congénital (PH, HCE) also funded this work. The human embryonic and fetal material was provided by the Joint MRC/Wellcome Trust (grant # 099175/Z/12/Z) Human Developmental Biology Resource (www.hdbr.org). None of the authors has a conflict of interest in relation to the work described herein.

